# Molecular mechanisms of mitochondrial Ca^2+^ exchanger NCLX

**DOI:** 10.64898/2026.01.28.702368

**Authors:** Li Zhang, Yan Han, Weizhong Zeng, Jing Xue, Yan Wang, Youxing Jiang

## Abstract

Mitochondrial Ca^2+^ homeostasis is maintained through coordinated influx and efflux processes, with NCLX long recognized as the primary Ca^2+^ extruder operating via Na^+^/Ca^2+^ exchange. Here, we report cryo-EM structures of rat NCLX in cytosolic-facing occluded and open states. The central transmembrane (TM) domain of NCLX comprises ten helices arranged in two inverted, structurally similar halves, with two α-repeats forming a central ion-binding pocket. Peripheral TMs 1 and 6 are loosely associated with the core and likely mediate alternative access to this site. These structural features closely resemble those of NCXs, indicating a conserved ion exchange mechanism. While NCLX retains the canonical Ca^2+^-binding site, it lacks several key Na^+^-binding residues found in NCXs, suggesting broader ion selectivity. Consistently, cell-based Ca^2+^ uptake assays show that NCLX mediates Ca^2+^ exchange using Na^+^, K^+^, Li^+^, and potentially protons as counterions. Based on the structural symmetry of NCLX and its bidirectional exchange capability, we propose a matrix-facing model and an alternating-access mechanism in which TMs 1 and 6 undergo sliding motions to enable ion exchange between cytosolic and matrix sides, analogous to NCX. These findings provide a structural and mechanistic framework for understanding NCLX-mediated Ca^2+^ transport in mitochondria.

## Introduction

Calcium is a critical secondary messenger in eukaryotic cells, regulating diverse processes ranging from metabolism to signal transduction^1-2^. Mitochondria serve as central hubs for calcium signaling, integrating metabolic demands with intracellular pathways^3-7^. Mitochondrial calcium concentrations are highly dynamic and tightly regulated^2,8^, with uptake primarily mediated by the mitochondrial calcium uniporter (MCU) complex, which facilitates calcium accumulation within the matrix^3,9-10^.

NCLX has long been recognized as the principal Na^+^/Ca^2+^ exchanger responsible for calcium efflux from mitochondria, a process essential for maintaining mitochondrial calcium homeostasis^8,11-12^. Localized to the inner mitochondrial membrane, NCLX mediates calcium extrusion in exchange for sodium^11-13^. Silencing NCLX strongly suppresses calcium efflux, whereas overexpression enhances it. Mutations, such as S258, disrupt mitochondrial calcium oscillations and impair downstream signaling^14^. Proper NCLX function is critical for mitochondrial health: knockout or dysfunction leads to Ca^2+^ overload, causing membrane depolarization, cristae disorganization, abnormal morphology, and impaired ATP synthesis and electron transport chain activity^15-16^. Beyond its mitochondrial functions, NCLX has also been implicated in several key physiological processes, including B lymphocyte calcium signaling, astrocyte-neuron communication, and cardiac energy regulation^17-21^. Notably, recent studies suggest that TMEM65, rather than NCLX, plays the determinant role in mitochondrial calcium regulation and export^22-25^, highlighting ongoing debate regarding its precise role.

Despite extensive functional studies, the molecular mechanism by which NCLX mediates calcium extrusion remains unclear, in part due to the challenges of manipulating the mitochondrial matrix environment. Here, we employ cryo-electron microscopy to determine the structures of rat NCLX in two conformational states, revealing conformational transitions associated with calcium release. Complementary cell-based Ca^2+^ uptake assays with plasma membrane-localized NCLX demonstrate that the exchanger mediates calcium transport in exchange for multiple monovalent cations, including Na^+^, K^+^, Li^+^, and potentially protons, establishing NCLX as a non-selective cation/Ca^2+^ exchanger.

## Results

### Overall structure of NCLX

To investigate the structure of NCLX, we expressed rat NCLX in HEK293S GnTI^−^ cells using the BacMam system and purified the protein in LMNG detergent. Size-exclusion chromatography (SEC) revealed two distinct elution peaks corresponding to a monomeric and a higher-molecular-weight oligomeric form, consistent with previous reports suggesting that mitochondrial NCLX can form oligomers (Supplementary Fig. 1a-b)^11^. Both fractions were subjected to single-particle cryo-EM analysis (Methods). Particles from the monomeric NCLX sample exhibited strong orientation bias, which was alleviated by adding fluorinated octyl maltoside (FOM) before grid freezing, yielding a reconstruction at 3.6 Å resolution (Supplementary Fig. 2 and Supplementary Table1). The larger SEC fraction contained trimeric NCLX, whose structure was determined at a resolution of 3.15 Å, with all three protomers adopting identical conformations (Fig. 1a, Supplementary Fig. 3 and Supplementary Table 1).

**Fig. 1:**
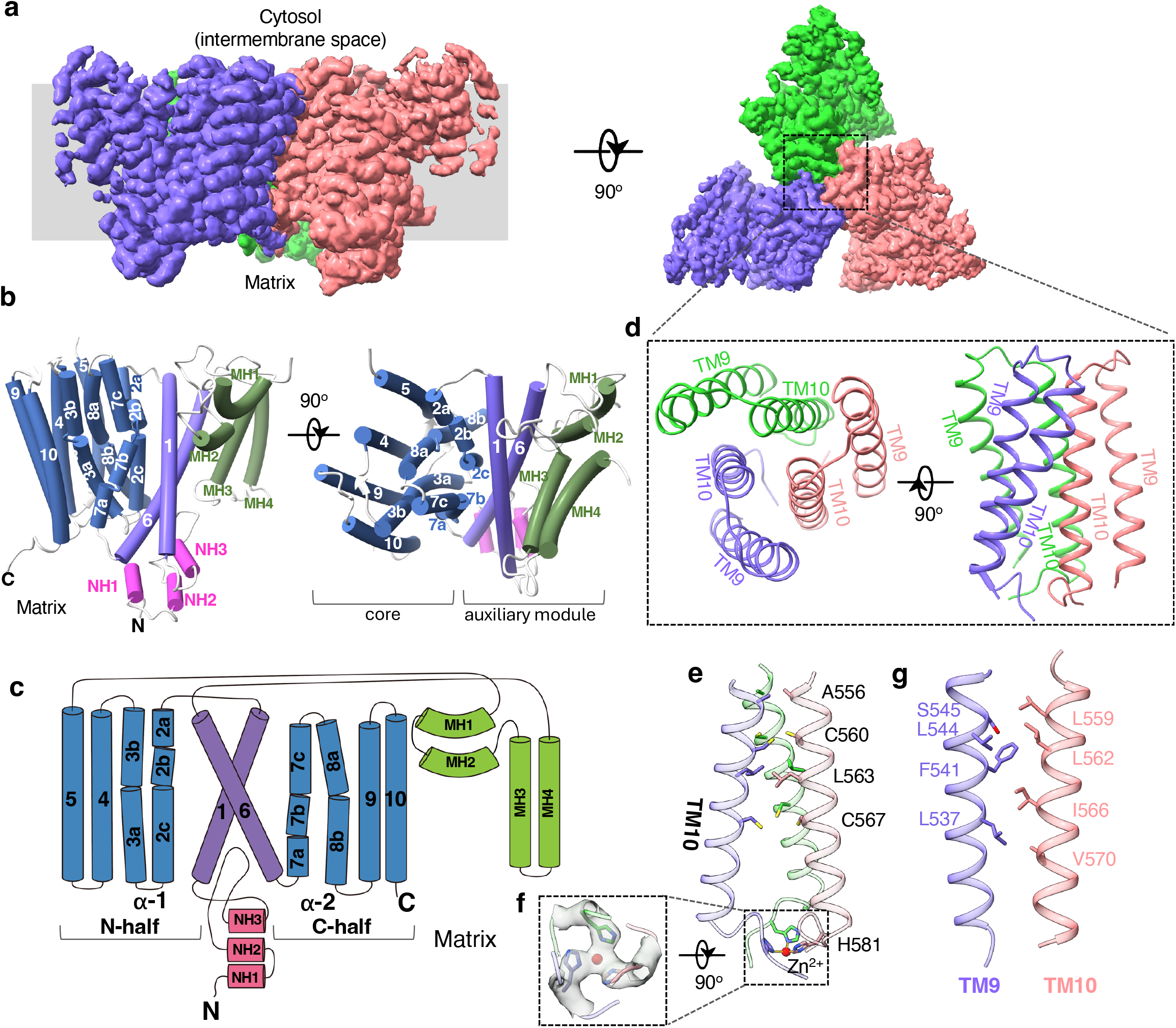
Overall structures of trimeric NCLX. **a** 3D reconstruction of the NCLX trimer shown in side and top views, with each subunit individually colored. **b** Cylinder representation of an NCLX subunit in side and top views. Transmembrane helices (TMs) 2-5 and 7-10 constitute the core of the exchanger, whereas TMs 1 and 6 together with the auxiliary helices (NH1-3 and MH1-4) form the auxiliary module that mediates alternating-access transitions. **c** Topology diagram of an NCLX subunit. **d** Top and side views of the trimerization assembly formed by TMs 9 and 10. **e** Zoomed-in view of the inter-subunit trimerization contacts within the TM10 helix bundle. **f** Zoomed-in view of the histidine cluster from the cytosolic side with density (grey surface) countered at 6σ. **g** Zoomed-in view of the inter-subunit contacts between TM10 and TM9 from a neighboring subunit.

Each NCLX protomer contains ten transmembrane (TM) helices arranged similarly to those of archaeal NCX (NCX_Mj) and the TM domain of human NCX1 (Fig. 1b-c and Supplementary Fig. 4) ^26-30^. In our structural comparison, the matrix-facing side of NCLX aligns with the extracellular side of NCXs (Supplementary Fig. 4). As in NCX, the ten TMs are organized into two pseudo-symmetrical halves (TM1-5 and TM6-10) with inverted topology (Fig. 1c). Eight helices (TMs 2-5 and 7-10) form a tightly packed core, containing two conserved α-repeats (TMs 2-3 as α-repeat I and TMs 7-8 as α-repeat II) that create a central ion-binding pocket essential for cation-calcium exchange (Fig. 1b). All α-repeat helices bend at their midpoints, defining the ion-binding site. TMs 1 and 6 are unusually long peripheral helices that loosely contact the core at ∼45° and likely move together as a “sliding-door”, mediating the alternating access transitions as seen in NCX exchangers (Fig. 1b).

In addition to the central TM domain, NCLX contains several auxiliary structural elements attached to TMs 1 and 6 (Fig. 1b-c). The N-terminal pre-TM1 region forms three short helices (NH1-3) lying on the luminal membrane surface, with the loop between NH2 and NH3 partially embedded in the membrane. The long TM5-TM6 linker forms two distinct helical modules. The first comprises two arch-shaped helices (MH1 and MH2) partially embedded in the cytosolic membrane leaflet. A similar structural element is also observed in human NCX1^26-28^. The second module consists of two antiparallel helices (MH3 and MH4) that insert deeply into but do not fully span the membrane. These helices are unique to NCLX and are best described as partially membrane-embedded elements rather than canonical TMs. Together, these auxiliary components form extensive hydrophobic contacts with TMs 1 and 6 and are separated from the TM core, except for a flexible loop that links them to TM5. This arrangement suggests that these elements likely move together with TMs 1 and 6 as a rigid body during conformational cycling.

Trimerization of NCLX is mediated primarily by TMs 9 and 10 (Fig. 1d-g). The N-terminal halves of the three TM10 helices bundle tightly around the threefold axis, forming extensive hydrophobic interactions (Fig. 1e). At their C-termini, three histidine residues (His581) cluster together and coordinate a strong non-protein density consistent with a metal ion (Fig. 1e-f). The geometry of this density resembles Zn^2+^ coordination by histidine observed in some zinc-binding proteins^31^. Given the abundance of transition metals - including Zn^2+^ - in the mitochondrial matrix^32^, we tentatively modeled this density as a Zn^2+^ ion without excluding the possibility of other transition metals. As further discussed later in the functional assays, this histidine-mediated interaction with a metal ion is important for stabilizing the trimeric assembly of NCLX. Each TM10 also interacts hydrophobically with TM9 from a neighboring subunit (Fig. 1g), further reinforcing trimer formation.

To assess the oligomeric state of NCLX in mitochondria, we also analyzed the size-exclusion chromatography profile of NCLX extracted from the mitochondria of NCLX-expressing cells (Supplementary Fig. 1c-d). NCLX isolated from mitochondria also eluted as two distinct peaks corresponding to trimer and monomer. Compared with NCLX purified from total cell extracts, the mitochondrial extract contained a higher proportion of the trimeric form. These results suggest that the trimeric form of NCLX exists in both mitochondrial and plasma membranes in our expression system. Given the tight inter-subunit trimerization interactions, we speculate that NCLX primarily forms trimers in its native state, and the monomer formation observed during purification likely results from detergent-induced disruption of these inter-subunit contacts. Further studies are required to verify the physiological relevance of NCLX trimerization. Because TMs 9 and 10 that mediate NCLX trimerization are spatially distant from the proposed gating module, trimerization is unlikely to influence the conformational changes associated with ion exchange.

### Ion binding sites

The four α-repeat helices enclose a central pocket deep within the membrane that serves as the substrate ion-binding site (Fig. 2a-b). Given the structural conservation between NCLX and NCX, the chemistry of NCLX ion binding pocket is compared with that of NCX_Mj whose substrate-binding sites are well defined (Fig. 2c)^29,30^. In NCX_Mj, the central pocket contains four substrate-binding sites: the central high-affinity Ca^2+^ binding S_Ca_ site, which can be occupied by Na^+^ in the absence of Ca^2+^; the intracellular and extracellular-facing S_int_ and S_ext_ sites, respectively, for Na^+^ binding; and the S_mid_ site right beside S_Ca_ for a water molecule. In comparison, only the S_Ca_ site is conserved in NCLX. The trimeric NCLX sample was prepared in a nominally calcium-free buffer without EGTA, resulting in a residual calcium concentration in the micromolar range. An EM density consistent with Ca^2+^ is observed at the position corresponding to the high-affinity Ca^2+^-binding site (S_Ca_) of NCX (Fig. 2b). The bound Ca^2+^ is coordinated by the carboxylate groups of Asp153 and Asp471 - acidic residues conserved between NCX and NCLX - as well as by the backbone carbonyls of Asn149 and Asn467 on TM2 and TM7, respectively.

**Fig. 2:**
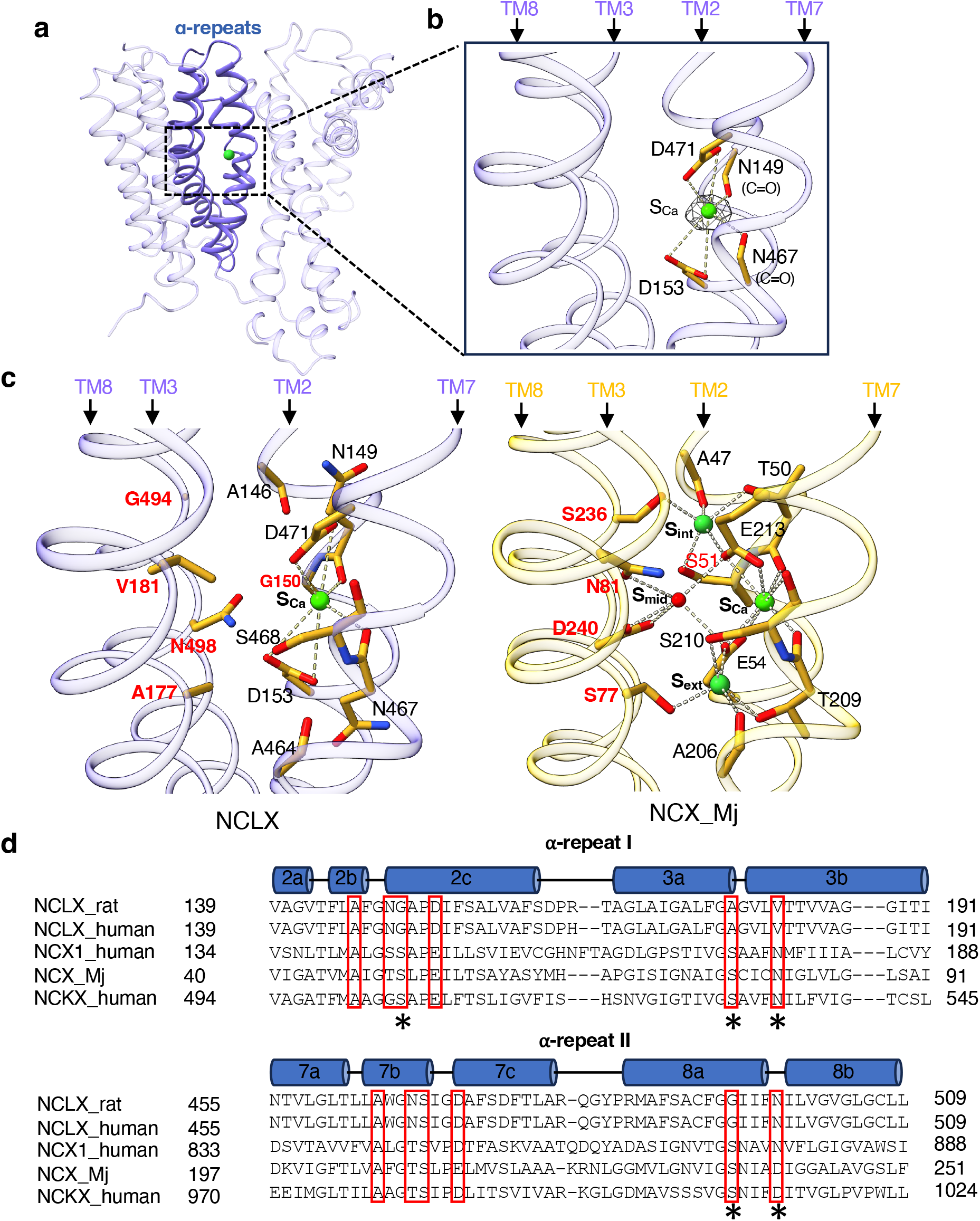
Ion-binding pocket of NCLX. **a** The two α-repeats (highlighted in blue) form the central ion-binding pocket of NCLX. **b** Zoomed-in view of the central pocket in trimeric NCLX, showing the density (grey mesh contoured at 6σ) at the S_ca_ site modeled as a Ca^2+^ ion. **c** Comparison of the central ion-binding sites between NCLX (left) and NCX_Mj (right; PDB 3V5U). Key residues that differ between NCLX and NCX_Mj at the S_int_, S_ext_ and S_mid_, sites are labeled in red. **d** Partial sequence alignment of rat NCLX (UniProt ID: Q6AXS0), human NCLX (UniProt ID: Q6J4K2), human NCX1 (UniProt ID: P32418), NCX_Mj (UniProt ID: Q57556), and human NCKX (UniProt ID: O06721) at the α-repeats. Residues forming the ion-binding sites in NCX are boxed in red. Asterisks indicate key residues that differ between NCX and NCLX within the ion-binding pocket.

In contrast, several highly conserved residues essential for Na^+^ binding in NCXs are absent in NCLX (Fig. 2c-d). When compared with NCX_Mj, Ser51 and Ser236 at the cytosolic-facing S_int_ site are replaced by Gly150 and Gly494, respectively, in NCLX; Ser77 at the S_ext_ site is replaced by Ala177 in NCLX; and Asn81 at the S_mid_ water-binding site is replaced by Val181. Substitution of these key Na^+^-chelating residues with smaller side chains enlarges the ion-binding pocket, creating a more open cavity. Together with the presence of the two acidic residues (Asp153 and Asp471) and several polar side chains, the ion-binding pocket of NCLX likely generates a negatively charged environment capable of accommodating multiple non-specific monovalent cations that serve as the exchanging ions for Ca^2+^.

### Transition from cytosolic-facing occluded to open state in NCLX

The core transmembrane (TM) region of trimeric NCLX (TMs 2-5 and 7-10) aligns closely with both the inward-facing human NCX1 and the outward-facing archaeal NCX_Mj structures (Supplementary Fig. 4), suggesting that this central core module remains largely static during the ion exchange cycle. In contrast, the peripheral TMs 1 and 6 of NCLX align well with those of NCX1 but differ from their counterparts in NCX_Mj, indicating that the NCLX structure captures a cytosolic-facing conformation analogous to the inward-facing state of NCX. During the transition to a matrix-facing conformation, TMs 1 and 6 of NCLX are likely to move to positions corresponding to their counterparts in NCX_Mj.

The monomeric NCLX structure is nearly identical to the trimeric form, except for a conformational change in the N-terminal segment of TM2 (TM2ab), which swings outward from the protein core (Fig. 3a). This displacement disrupts the Ca^2+^-chelating backbone carbonyl of Asn149 from the S_Ca_ site (Fig. 3a inset). Notably, the monomeric NCLX sample was prepared in the presence of 1 mM EGTA and high NaCl, yet an ion density remains at the S_Ca_ site (Fig. 3b). Given that S_Ca_ serves as a shared binding site for both Ca^2+^ and Na^+^ in NCX, this density most likely corresponds to Na^+^ in our sample conditions.

**Fig. 3:**
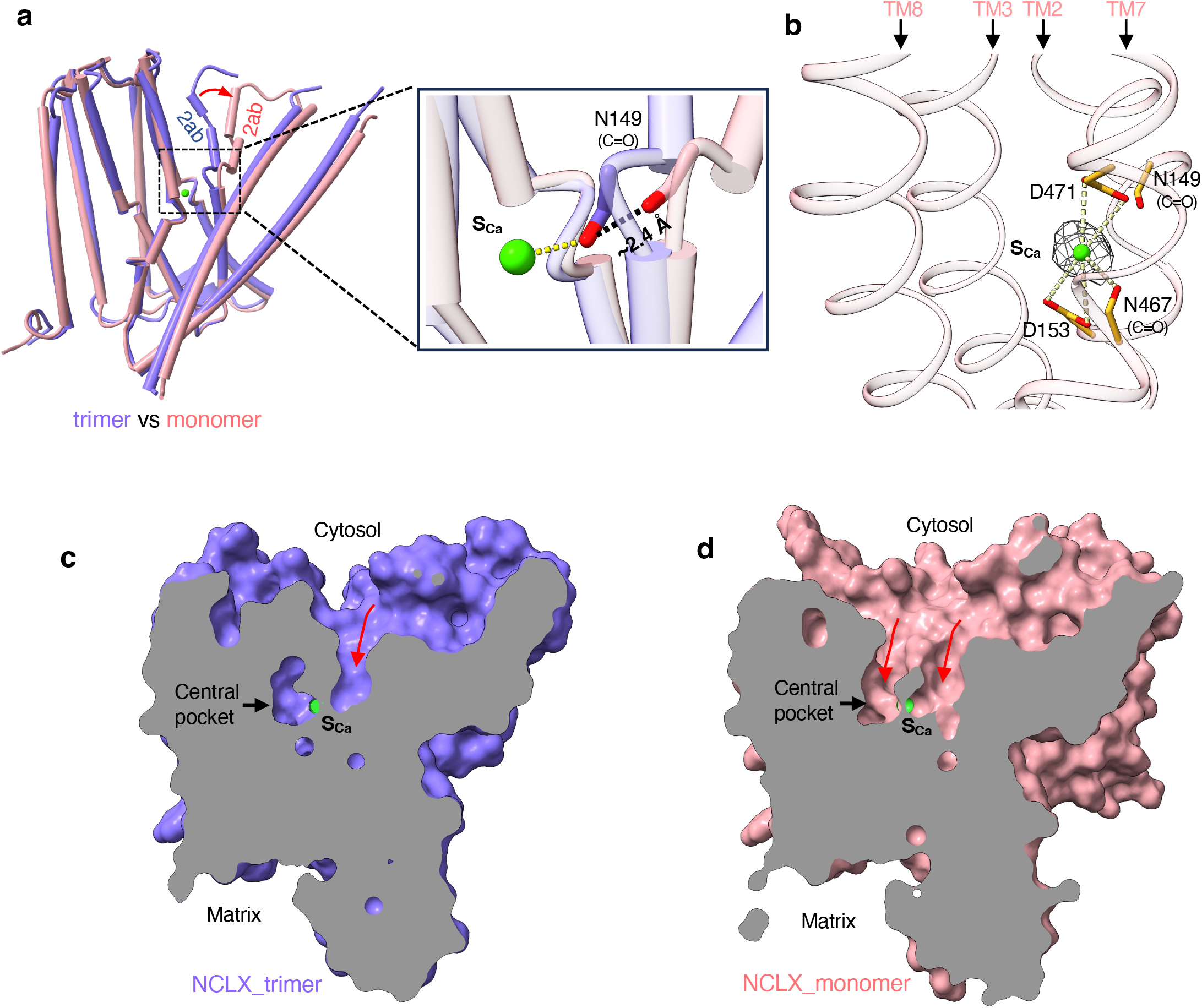
Transition of cytosolic-facing NCLX from the occluded to the open state. **a** Structural comparison between trimeric (blue) and monomeric (pink) NCLX. The red arrow highlights the conformational change of TM2ab. For clarity, TM5 is omitted in this view. Inset: the backbone carbonyl of N149 shifts away from the S_Ca_ site. **b** Zoomed-in view of the central pocket of monomeric NCLX, showing the density (grey mesh contoured at 6σ) at the S_Ca_ site modeled as a Na^+^ ion. **c** Cross-section of the surface-rendered trimeric NCLX structure showing a deep cavity (red arrow) extending toward the S_Ca_ site, while the central ion-binding pocket remains occluded. **d** Cross-section of the surface-rendered monomeric NCLX structure showing that both the S_Ca_ site and the central ion-binding pocket are exposed to the cytosol through two independent solvent-accessible pathways (red arrows).

In the trimeric NCLX structure, a pronounced cavity is observed on the cytosolic surface, extending deep toward the Ca^2+^-binding site (Fig. 3c). Despite this, the central ion-binding pocket remains occluded from the cytosol, indicating that the trimeric structure represents a cytosolic-facing occluded state. In the monomeric NCLX, the swing movement of TM2ab creates a continuous, cytosol-facing ion permeation pathway, rendering the central pocket solvent-accessible (Fig. 3d). Thus, the structural differences between trimeric and monomeric NCLX reflect a transition from a cytosolic-facing occluded state to a cytosolic-open state. The emergence of two independent solvent-accessible pathways for ion access in the open NCLX closely parallels the two ion pathways observed in the structures of outward-facing NCX_Mj^29-30^. Moreover, the key conformational change between the open and occluded states occurs at equivalent regions of the symmetry-related α-repeat helices - TM2ab in NCLX and TM7ab in NCX_Mj, highlighting a conserved structural mechanism underlying ion exchange in NCX and NCLX

### Alternative access mechanism

Although the resolved NCLX structures adopt cytosolic-facing conformations, the close structural similarity between NCLX and NCX, together with the internal symmetry of its TM module, suggests that a matrix-facing conformation would preserve the same overall architecture and symmetry. In this alternate state, the ion access pathways would be reoriented toward the matrix side, effectively forming an inverted counterpart of the cytosolic-facing structure. As depicted in Supplementary Fig. 5, superimposing the 10-TM module of the trimeric NCLX with its inverted copy yields a simple and mechanistically consistent model for the matrix-facing conformation. This superimposition shows near-perfect alignment of the central α-repeat helix bundle, while the peripheral helices TM1 and TM6 occupy distinct positions. By translocating TM1 and TM6, together with their associated auxiliary helices (NH1-3 and MH1-4), which likely move as a rigid unit, to the positions corresponding to the inverted TM6 and TM1, respectively, we can generate a plausible matrix-facing occluded model (Supplementary Fig. 5). Similar symmetry-based approaches have been used to construct alternate-state models for NCX^26^ and other secondary solute transporters containing inverted structural repeats^33-34^. Our cytosolic-facing structure and predicted matrix-facing conformation closely align with recently reported NCLX structures in both cytosolic- and matrix-facing states (Supplementary Fig. 5d-e)^35^.

Compared with the cytosolic-facing occluded structure, the matrix-facing model suggests that TMs 1 and 6 undergo a coordinated “sliding” motion between the two conformations (Fig. 4a), consistent with the alternating-access mechanism established for NCXs. From this matrix-facing occluded model, we further infer that TM7ab would undergo a conformational swing analogous to that observed in TM2ab of the monomeric NCLX, opening the central ion-binding pocket to the matrix side (Fig. 4b). Together, the trimeric and monomeric NCLX structures, combined with the predicted matrix-facing model, support a simple alternating-access mechanism for NCLX ion exchange, as summarized in Fig. 4b.

**Fig. 4:**
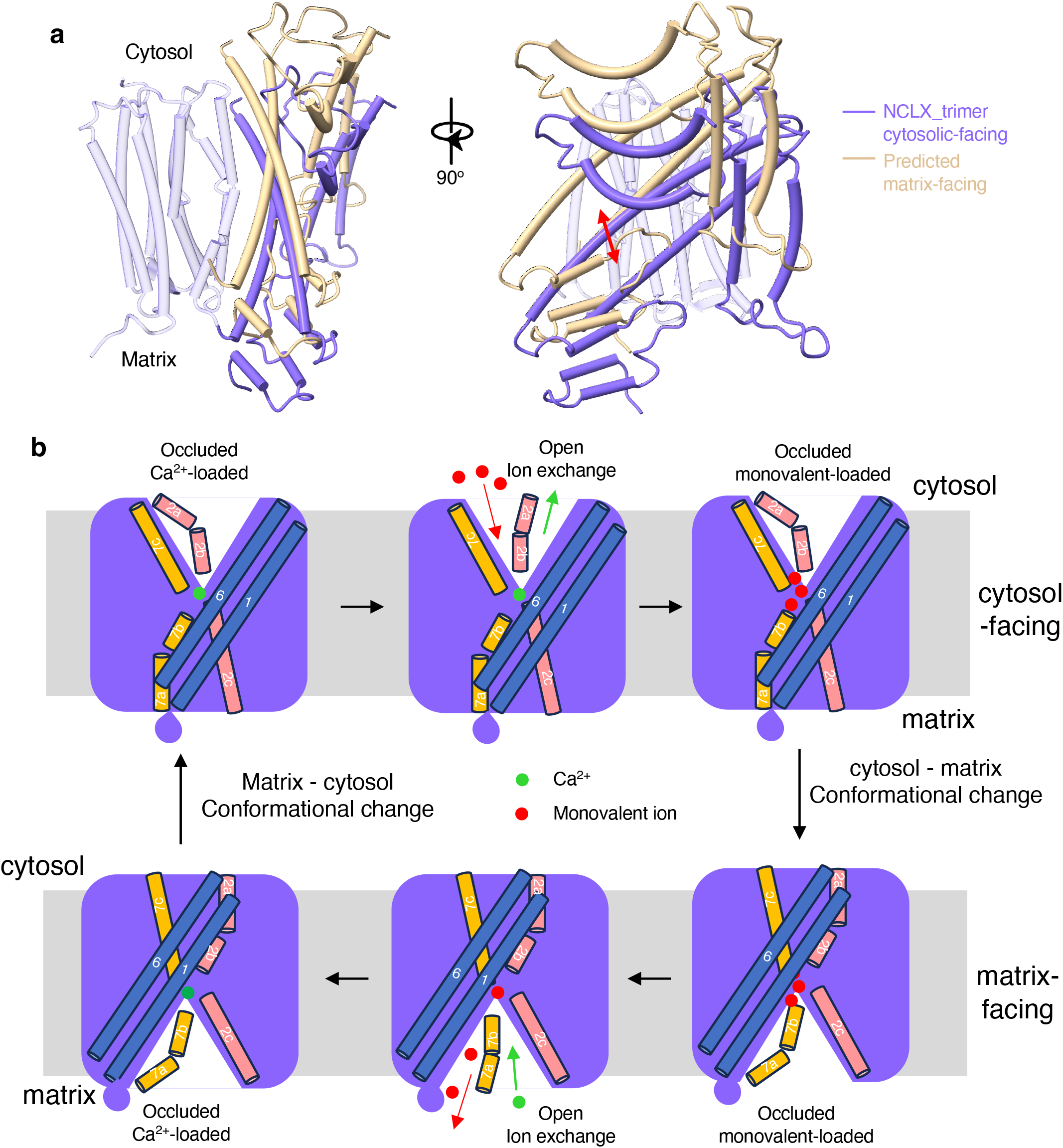
Alternative access mechanism of NCLX. **a** Structural comparison between the cytosolic-facing NCLX from the trimer (blue) and the proposed matrix-facing NCLX (light gold). The core regions (light blue) remain unchanged between the two conformations. Tms 1 and 6, together with their associated auxiliary components, undergo a rigid-body sliding motion, as indicated by the red arrow. **b** Cartoon representation of the proposed alternating-access cycle of cation/Ca^2+^ exchange in NCLX. Although multiple monovalent cations may be required to exchange a single Ca^2+^ ion, the exact stoichiometry between monovalent cations and Ca^2+^ remains unresolved.

### NCLX functions as a slow, non-selective cation/Ca^2+^ exchanger

The high structural similarity between NCLX and NCX, together with the conserved central Ca^2+^-binding site, suggests that NCLX likely mediates Ca^2+^ exchange through an alternating-access mechanism involving cytosolic-matrix conformational transitions. However, the absence of several key Na^+^-binding residues in the α-repeats of NCLX raises the possibility that it functions as a non-selective cation/Ca^2+^ exchanger rather than a strict Na^+^/Ca^2+^ exchanger. To test this hypothesis, we performed cell-based Ca^2+^ uptake assays using HEK cells expressing rat NCLX (Methods). Plasma membrane localization of overexpressed NCLX was confirmed by cell-surface biotinylation (Supplementary Fig. 6a and Methods). Cells transfected with an empty vector served as negative controls, while cells expressing human NCX1 were used as positive controls for Ca^2+^ uptake. All transfected cells were loaded with the Ca^2+^-sensitive dye fura-2, and cytosolic [Ca^2+^] changes were monitored by the fluorescence ratio at 510 nm (excitation 340/380 nm) (Fig. 5, Supplementary Fig. 6-c, and Methods)^29,36^.

**Fig. 5:**
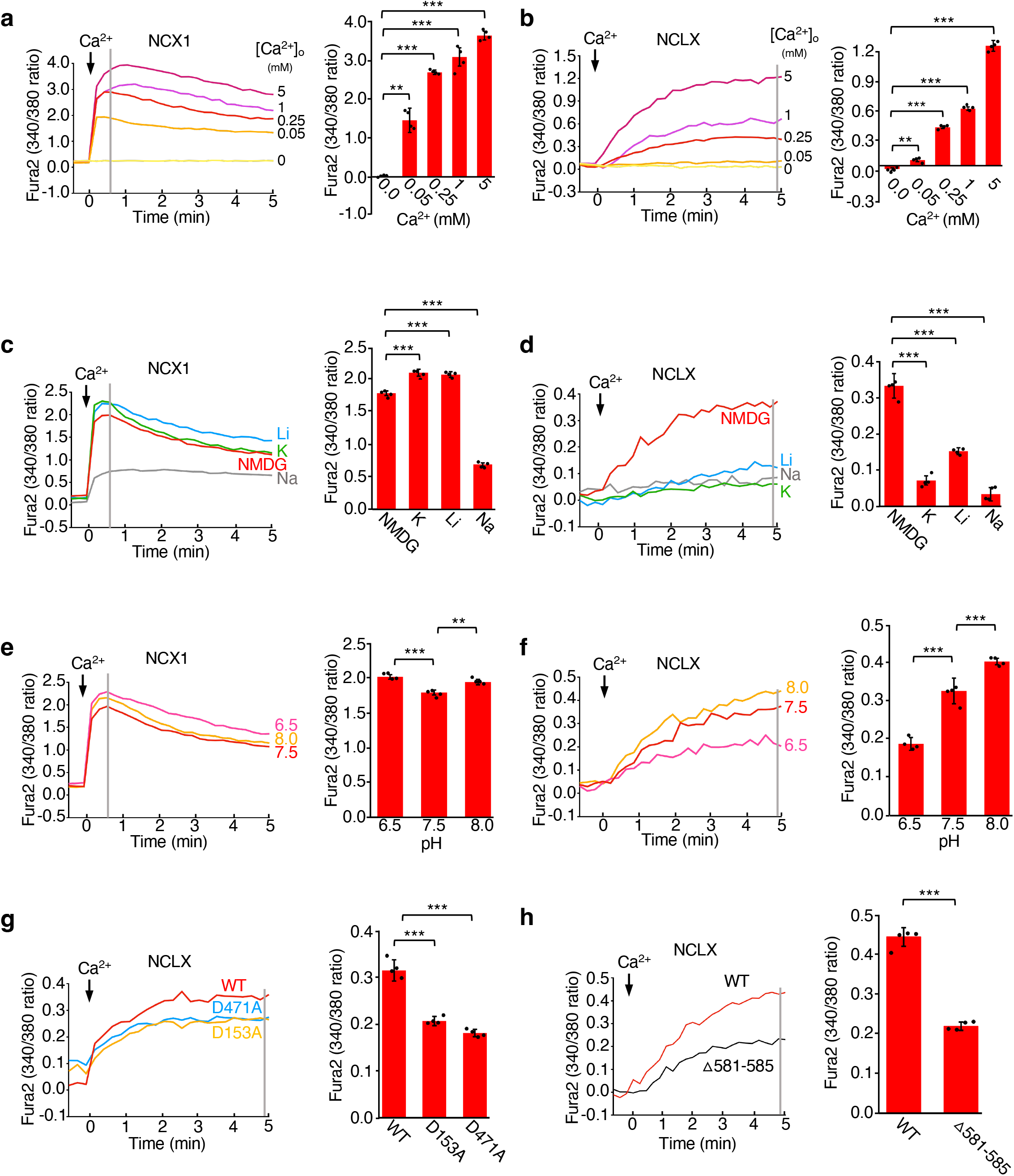
Ca^2+^ uptake assays in exchanger-expressing HEK cells. **a** Sample traces of Ca^2+^ uptake in NCX1-expressing HEK cells using 150 mM NMDG^+^ as the monovalent salt and varying external [Ca^2+^] (left panel). Cytosolic [Ca^2+^] changes were monitored by fura-2 fluorescence ratios at 510 nm (excitation 340/380 nm) following the addition of external Ca^2+^ (indicated by the arrow) and were quantified at the time point marked by the grey line (right panel). All data points shown here and the subsequent experiments represent mean ± SEM of n = 4 independent replicates. **b** Same experiment as in **(a)** performed with NCLX-expressing HEK cells. All traces in **(a)** and **(b)** are shown after background subtraction (Supplementary Fig. 6). **c** Ca^2+^ uptake in NCX1-expressing HEK cells with 0.05 mM external [Ca^2+^] and 150 mM of various monovalent salts. **d** Ca^2+^ uptake in NCLX-expressing HEK cells with 0.25 mM external [Ca^2+^] and 150 mM of various monovalent salts. **e** Ca^2+^ uptake in NCX1-expressing HEK cells with 0.05 mM [Ca^2+^] and 150 mM NMDG^+^ at different extracellular pHs (6.5, 7.5, and 8.0). **f** Ca^2+^ uptake in NCLX-expressing HEK cells with 0.25 mM [Ca^2+^] and 150 mM NMDG^+^ at different extracellular pHs (6.5, 7.5, and 8.0). **g, h** Ca^2+^ uptake in NCLX WT or mutant-expressing HEK cells with 0.25 mM external [Ca^2+^] and 150 mM NMDG^+^ as the monovalent salt. Statistical analysis was performed using a two-sided paired t-test. Source data are provided as a Source Data file. *** represents p < 0.001 and ** represents p < 0.01.

In the first experiment, we measured exchanger-mediated Ca^2+^ uptake at various extracellular [Ca^2+^] using 150 mM NMDG^+^ as the monovalent salt (Fig. 5a-b). NCX1-expressing cells displayed a rapid cytosolic Ca^2+^ increase in a concentration-dependent manner, which peaked within one minute and then declined gradually, likely due to mitochondrial Ca^2+^ uptake via the MCU pathway (Fig. 5a and Supplementary Fig. 6b). NCLX-expressing cells also exhibited a cytosolic [Ca^2+^] rise upon external Ca^2+^ addition, but with a markedly slower rate and a lower steady-state cytosolic [Ca^2+^] (Fig. 5b and Supplementary Fig. 6c). These results demonstrate that NCLX functions as a Ca^2+^ exchanger, similar to NCX1, but with substantially slower transport kinetics. Because NCX1 utilizes intracellular Na^+^ as the counterion for Ca^2+^ influx, extracellular Na^+^, but not other monovalent cations, can effectively compete against Ca^2+^ binding and inhibit NCX1-mediated Ca^2+^ uptake. In contrast, if NCLX acts as a non-selective cation/Ca^2+^ exchanger, multiple monovalent cations could serve as alternative counterions, and their presence in the extracellular solution should also inhibit Ca^2+^ uptake. To test this, we performed a second set of uptake assays at fixed Ca^2+^ concentrations (0.05 mM for NCX1; 0.25 mM for NCLX) using 150 mM of various monovalent salts (XCl, where X = Li^+^, Na^+^, K^+^, or NMDG^+^) (Fig. 5c-d). Consistent with expectations, extracellular Na^+^ strongly inhibited NCX1-mediated Ca^2+^ uptake compared to NMDG^+^, whereas Li^+^ and K^+^ had no inhibitory effect - in fact, they slightly enhanced Ca^2+^ entry (Fig. 5c). In contrast, all three monovalent cations (Li^+^, Na^+^, and K^+^) significantly suppressed NCLX-mediated Ca^2+^ uptake, supporting the hypothesis that NCLX functions as a non-selective cation/Ca^2+^ exchanger (Fig. 5d).

Given recent reports suggesting that NCLX may also operate as a proton/Ca^2+^ exchanger in mitochondria^35^, we further tested the pH dependence of Ca^2+^ uptake at three extracellular pH values (6.5, 7.5, and 8.0) using NMDG^+^ as the monovalent ion. While NCX1-mediated uptake was largely pH-insensitive (Fig. 5e), NCLX-mediated Ca^2+^ uptake progressively increased at higher pH (Fig. 5f), raising the possibility that protons, likely in the form of hydronium ions (H_3_O^+^), can also serve as exchangeable cations for NCLX.

We further examined the functional roles of two key structural elements of NCLX. First, the conserved acidic residues Asp153 and Asp471 at the central Ca^2+^-binding site were individually mutated to alanine. Although these mutants exhibited expression levels comparable to wild-type NCLX (Supplementary Fig. 7a), they showed markedly reduced Ca^2+^ uptake, confirming the essential role of these residues in Ca^2+^ binding and transport (Fig. 5g). Second, we generated a deletion mutant (Δ581-585) lacking the C-terminal tail (residues 581-585) involved in His581-mediated metal coordination and trimerization. With comparable expression levels, size-exclusion chromatography revealed a substantial reduction in trimer formation (Supplementary Fig. 7b-d). Functionally, this mutant also exhibited significantly decreased Ca^2+^ uptake relative to wild-type NCLX (Fig. 5h). Together, these results indicate that the C-terminal tail is critical for stabilizing trimer assembly and that trimerization enhances NCLX exchange activity.

## Discussion

In this study, we establish the structural and functional basis of mammalian NCLX. The trimeric NCLX structure captures a cytosolic-facing occluded state, whereas the monomeric structure represents a cytosolic-facing open state. Although the native oligomeric state of NCLX in mitochondria remains to be determined, the extensive inter-subunit interactions observed in our structure suggest that NCLX predominantly assembles as a trimer *in vivo*. The central transmembrane module of NCLX adopts the same architectural framework as NCX family exchangers, underscoring a conserved structural mechanism for ion exchange. Leveraging the internal symmetry and bidirectional transport property of NCLX, we also generated a structural model for the matrix-facing conformation. Together, these structures and models support an alternating-access mechanism driven by the sliding movement of TMs 1 and 6, analogous to that in NCX. Trimeric NCLX structures in multiple conformations were also recently reported by Fan *et al*^35^. Our cytosolic-facing trimer closely matches their structure (Supplementary Fig. 5e), and both studies identify a similar Ca^2+^-dependent conformational change at TM2ab. In addition, Fan *et al*^35^. resolved a matrix-facing NCLX structure, providing direct insight into the conformational transitions underlying alternating ion access during the Ca^2+^ exchange cycle. Notably, our symmetry-based model of the matrix-facing state aligns closely with their experimentally determined structure, recapitulating the same inward-outward conformational rearrangement.

NCLX retains a conserved Ca^2+^-binding site but lacks several Na^+^-coordinating residues present in canonical NCXs, suggesting that it functions as a non-selective cation/Ca^2+^ exchanger rather than a strict Na^+^/Ca^2+^ exchanger. Consistent with this idea, our cell-based assays show that NCLX-mediated Ca^2+^ uptake is suppressed by multiple extracellular monovalent cations, including Na^+^, K^+^, and Li^+^, indicating that these ions can serve as counterions for Ca^2+^ transport (Fig. 5d). Although such competition assays cannot fully exclude the possibility that these monovalent ions act as blockers rather than participating directly in ion exchange, several observations support their role as counter-exchanging ions. First, Ca^2+^ uptake is not completely abolished even at high concentrations of monovalent cations, arguing against simple competitive blockade and instead suggesting that reduced uptake reflects an increased inward gradient of counterions opposing Ca^2+^ influx. Second, in Na^+^-selective NCX1, only Na^+^ suppresses Ca^2+^ uptake, whereas in NCLX, Li^+^, Na^+^, and K^+^ produce comparable inhibitory effects. This makes it unlikely that Na^+^ acts as the exchange ion in NCX1 while these same monovalent cations function as blockers in NCLX. Third, the high structural similarity between NCX1 and NCLX, particularly within the α-repeat regions that mediate ion exchange, supports a conserved alternating-access mechanism. The absence of specific Na^+^-binding residues in NCLX likely permits broader access of monovalent cations to the ion-binding pocket. Within this framework, monovalent cations that enter the pocket from one side at high concentration are expected to be released to the opposite side during conformational cycling, thereby functioning as counterions in Ca^2+^ transport.

The primary discrepancy between our study and that of Fan *et al* lies in the functional characterization of NCLX^35^. While their results support exclusive H^+^/Ca^2+^ exchange activity, our assays indicate that NCLX operates as a non-selective cation/Ca^2+^ exchanger, capable of coupling Ca^2+^ transport to Na^+^, K^+^, Li^+^, and potentially H^+^. Given the distinct experimental approaches used in the two studies, this difference is not easily reconciled. Notably, all of our functional assays were conducted alongside NCX1, a well-established Na^+^-selective Ca^2+^ exchanger, under identical conditions. Under these conditions, NCLX-mediated Ca^2+^ transport is modulated by multiple monovalent cations (Li^+^, Na^+^, and K^+^) as well as by protons, whereas NCX1 activity is selectively affected by Na^+^ alone. These comparative results support a broader ion selectivity for NCLX.

The stoichiometry proposed in our working model (Fig. 4b) is inferred from previously characterized Na^+^/Ca^2+^ exchangers, including the extensively studied NCX1 and the high-resolution structure of the archaeal homolog NCX_Mj, both of which share conserved structural features with NCLX. A 3:1 Na^+^/Ca^2+^ exchange ratio has been well established for NCX through electrophysiological measurements^37^. In NCX, high-affinity Ca^2+^ binding at the S_Ca_ site is progressively weakened by the binding of multiple Na^+^ ions, which reduce Ca^2+^ affinity through charge neutralization and electrostatic repulsion. This process ultimately drives Ca^2+^ release during the exchange cycle. Given the strong structural conservation of the ion-binding pocket between NCX and NCLX, we propose that NCLX likely adopts a similar 3:1 stoichiometry for monovalent cation/Ca^2+^ exchange.

Several recent studies have identified TMEM65, another inner mitochondrial membrane protein, as a key contributor to mitochondrial Ca^2+^ regulation and efflux, either directly or through modulation of NCLX^22-25^. In one study, TMEM65 has been reported to interact with NCLX and regulate mitochondrial Ca^2+^ efflux in an NCLX-dependent manner^22^. In contrast, two other studies have proposed that TMEM65, rather than NCLX, directly mediates rapid mitochondrial Ca^2+^ extrusion^23,25^. Notably, TMEM65 exhibits a more restricted expression pattern, enriched in excitable tissues, whereas NCLX is broadly expressed^23,38^. Although our study does not resolve this discrepancy, our structural and functional data demonstrate that NCLX possesses intrinsic Ca^2+^ exchange activity and likely remains an essential regulator of mitochondrial Ca^2+^ homeostasis.

## Methods

### Protein expression and purification of NCLX

Rat NCLX cDNA was synthesized and codon-optimized for mammalian expression (GenScript). The cDNAs were subcloned into the pEZT-BM vector with a C-terminal Strep tag. NCLX was expressed in HEK GnTI^−^ cells using the BacMam system. Bacmids were generated in *E. coli* DH10Bac cells, and baculovirus was produced in Sf9 cells. For protein expression, the baculovirus carrying the NCLX gene was used to infect HEK GnTI^−^ cells at a 1:50 (v/v) ratio. Cultures were maintained at 37 °C, 125 rpm, and 8% CO_2_. Twelve hours after infection, 10 mM sodium butyrate was added to enhance protein expression, and the cells were cultured at 30 °C, 125 rpm, and 8% CO_2_ for 60 hours before harvesting by centrifugation at 3,500 × *g* for 15 minutes. The same protocol was used for the expression of NCLX D153A, D471A, and the C-terminal truncation (Δ581-585) mutants.

The collected cells were resuspended in lysis buffer containing 150 mM NaCl, 25 mM HEPES (pH 7.4), 2 μg/ml DNase I, 0.5 μg/ml pepstatin, 2 μg/ml leupeptin, and 1 μg/ml aprotinin. Cells were disrupted by sonication, and membrane proteins were extracted with 1% LMNG at 4 °C for 2 hours. The supernatant was collected after centrifugation at 45,000 × *g* for 45 minutes, and was incubated with Strep-Tactin XT resin for 1 hour at 4 °C. The resin was collected by centrifugation at 500 × *g* for 5 minutes and washed with 20 column volumes of wash buffer (150 mM NaCl, 25 mM HEPES pH 7.4, 0.03% LMNG). Bound protein was eluted with wash buffer supplemented with 50 mM biotin.

For further purification, the eluted protein was concentrated and subjected to size-exclusion chromatography using a Superdex 200 Increase 10/300 GL column (Cytiva) equilibrated with the wash buffer. The peak fractions were collected and concentrated to approximately 2 mg/ml for cryo-EM grid preparation. For the purification of rat monomeric NCLX, 1 mM EGTA was added to the mobile phase during SEC.

### Cryo-EM sample preparation and data acquisition

Cryo-EM grids were prepared using a Mark IV Vitrobot (Thermo Fisher) with the chamber maintained at 100% humidity and 8°C. Fresh protein sample (4 µL) was applied to glow-discharged Quantifoil R1.2/1.3 300-mesh gold holey-carbon grids (Quantifoil). After incubation for 5 s, the grids were blotted with a blot force of 8 for 3 s and immediately plunged into liquid ethane precooled by liquid nitrogen. For preparation of rat monomeric NCLX grids in the presence of FOM, 0.7 mM FOM was added to the protein sample, briefly mixed, and subjected to the same vitrification procedure as described above.

Sample grids of monomeric NCLX, with or without FOM, were imaged on a 300 kV Titan Krios microscope (Thermo Fisher) equipped with a Falcon 4i detector. Movie stacks were recorded using SerialEM at a magnification of 165,000×. Data were recorded with a defocus range of −1.1 to −2.5 µm, a total exposure time of 4 s, and a cumulative electron dose of 60 e^−^/Å^2^. The calibrated pixel size for the dataset was 0.743 Å.

For trimeric NCLX, data collection was performed on a 300 kV Titan Krios (Thermo Fisher) equipped with a K3 Summit direct electron detector (Gatan). A GIF Quantum energy filter with a 20 eV slit width was applied during acquisition. Images were collected in SerialEM at a magnification of 130,000×, corresponding to a calibrated super-resolution pixel size of 0.2555 Å. Data were acquired with a defocus range of −1.1 to −2.5 µm and a total accumulated electron dose of 60 e^−^/Å^2^.

### Cryo-EM data processing

For the rat monomeric NCLX dataset, 6,720 movie stacks from samples without FOM were collected, motion corrected with RELION, and CTF parameters estimated using GCTF35-36. Gautomatch (Kai Zhang, https://www.mrc-lmb.cam.ac.uk/kzhang/) particle picking yielded 2,325,645 particles. Similarly, 7,326 stacks from samples with FOM produced 2,290,823 particles. All particles were extracted with threefold binning, combined, and subjected to 3D classification. A subset of 467,738 particles was re-extracted without binning and refined. A tight mask encompassing the rigid body of the protein was subsequently applied for further 3D classification, resulting in 336,224 particles. These particles were refined, followed by two rounds of CTF refinement and post-processing, producing a 3.62 Å density map. Resolution was estimated by the gold-standard Fourier shell correlation (FSC) at the 0.143 criterion, and local resolution was assessed in RELION.

For the rat trimeric NCLX dataset, 7,290 movie stacks were acquired and motion corrected with MotionCor2, and CTF parameters were estimated using GCTF^39-40^. Gautomatch (Kai Zhang, https://www.mrc-lmb.cam.ac.uk/kzhang/) particle picking yielded 1,360,717 particles, which were extracted with eightfold binning and subjected to 2D classification. A total of 1,131,535 particles were selected for initial 3D classification without symmetry, after which 349,174 particles were re-extracted with fourfold binning and refined with C3 symmetry. Subsequent 3D classification with C3 symmetry yielded 205,993 particles, which were refined and further classified with a tight mask around the rigid body of the protein. This resulted in 47,504 particles, which were subjected to 3D refinement followed by CTF refinement. During CTF refinement, microscope-induced aberrations were first corrected at the micrograph level, including beam tilt, trefoil, and 4^th^ order aberrations, along with the estimation of anisotropic magnification. Subsequently, per-particle defocus values were refined to account for local variations across individual particles. Two cycles of these CTF refinements were performed to ensure the parameters’ convergence. Finally, post-processing was performed with a mask covering the protein density to improve the interpretability of the cryo-EM density map and enhance high-resolution features by B-factor sharpening, producing a 3.15 Å density map. Resolution was determined by the gold-standard Fourier shell correlation (FSC) at the 0.143 criterion, and local resolution was assessed in RELION. The final map was further improved with DeepEMhancer for model building^41^.

### Model building and refinement

The initial structural model of NCLX was generated using ModelAngelo^42^. For detailed modeling, the initial model was fitted into the density map and manually adjusted in Coot^43^. The rebuilt model was further refined using the real_space_refine program in PHENIX^44^. All structures were validated by assessing MolProbity scores and Ramachandran plot statistics. Structural figures were prepared using PyMOL (Schrödinger, LLC), Chimera, and ChimeraX^45-46^.

### Cell-based calcium uptake assay

A similar protocol as previously described was used to detect the cell-based calcium uptake assay^29^. GnTI^-^ cells were infected at a density of 2.5 × 10^6^ cells/mL with baculovirus carrying the rat NCLX gene, human NCX1 gene, or an empty vector at a 1:40 (v/v) ratio. Cultures were maintained at 37 °C, 125 rpm, and 8% CO_2_. Twelve hours post-infection, 10 mM sodium butyrate was added to enhance protein expression, and the culture was shifted to 30 °C, 125 rpm, and 8% CO_2_ for further culture. NCLX-infected cells and their corresponding controls were cultured for a total of 60 hours after sodium butyrate addition, whereas NCX1-infected cells and their controls were cultured for 48 hours.

After the expression period, 10 mL of cells were harvested by centrifugation (200 × g, 2 min), washed with fresh medium, and resuspended in 1 mL. Cells were then loaded with 7.5 μM Fura-2 AM at 30 °C for 30 min. Following dye loading, cells were pelleted (500 × g, 1 min), washed twice, and resuspended in assay buffer containing 150 mM XCl, 5 mM KCl, 1 mM MgCl_2_, 0.5 mM EGTA, 10 mM Tris-HCl (pH 7.5), 10 mM glucose, and 0.1% BSA, where X was NMDG, Li^+^, Na^+^ or K^+^.

For the proton selectivity assay, Tris-HCl buffer at pH 8.0 or MES buffer at pH 6.5 was used in place of the standard Tris-HCl buffer at pH 7.5. Fluorescence measurements were performed on a SpectraMax M6 spectrophotofluorometer (Molecular Devices) at 510 nm emission with dual excitation at 340 nm and 380 nm. The 340/380 nm ratio (R) was used to monitor changes in intracellular Ca^2+^ concentration.

### Cell-surface protein isolation and western blot

For the cell-surface protein biotinylation and isolation, they were performed with the cell surface protein biotinylation and isolation kit (Pierce, A44390) following the manufacturer’s instructions. Briefly, 10 mL infected cells were harvested by centrifugation (200 × g, 2 min) and washed twice with PBS. For surface labeling, cells were resuspended in PBS containing Sulfo-NHS-SS-Biotin and incubated at room temperature for 10 min with gentle mixing. The reaction was stopped by centrifugation and two washes with ice-cold TBS provided in the kit. Hereafter, cells were lysed withthe kit lysis buffer, and clarified lysates were incubated with NeutrAvidin™ Agarose to enrich biotinylated proteins. Bound proteins were eluted with 10 mM DTT, separated by SDS-PAGE, and transferred to PVDF membranes for immunoblotting. Anti-NCLX (Invitrogen, PA5-114330) was used to detect cell-surface NCLX. In parallel, prior to cell-surface protein labeling, 500 µL of cell suspension from each sample was collected by centrifugation and lysed with RIPA buffer (Pierce, 89900) to prepare total lysates. These lysates were probed with anti-GAPDH (Invitrogen, 39-8600) as an input control.

### Mitochondria isolation

Mitochondria were isolated using a protocol similar to that previously reported^47^. Briefly, 8 L of NCLX-expressing GnTI^-^ cells at a density of 2.0 × 10^6^ cells/mL were harvested by centrifugation at 600 × g for 10 min and washed twice with PBS. After determining the pellet weight, the cells were resuspended in hypotonic buffer containing 20 mM KCl, 0.5 mM MgCl2, and 10 mM Tris-HCl (pH 7.4) at a ratio of 1 mL buffer per 0.15 g of cells. The suspension was incubated on ice for 5 min to allow cell swelling and then transferred to a pre-cooled Teflon-glass Dounce homogenizer, followed by 25 times up-and-down passing the pestle. The homogenate was adjusted to isotonic conditions by adding 1 M sucrose buffer (1 M sucrose, 10 mM Tris-HCl, pH 7.4) at a ratio of 1:3 (v/v). The lysate was centrifuged at 1,200 × g for 10 min, and the supernatant was transferred to a new tube. This centrifugation step was repeated until no visible pellet remained. The pooled supernatant was then centrifuged at 150,000 × g for 5 min to collect the mitochondrial fraction. The resulting pellet was washed with buffer containing 250 mM sucrose and 10 mM Tris-HCl (pH 7.4). Finally, the pellet was flash-frozen in liquid nitrogen and stored at −80°C until further use.

## Acknowledgement

Single particle cryo-EM data were collected at the University of Texas Southwestern Medical Center Cryo-EM Facility (CEMF) funded by the CPRIT Core Facility Support Award RP170644, at Pacific Northwest Center for Cryo-EM for structural studies supported by proposal 160365, and at Howard Hughes Medical Institute Janelia Cryo-EM Facility. Cryo-EM sample grids were prepared at the Structural Biology Laboratory at UT Southwestern Medical Center, partially supported by grant RP170644 from CPRIT. This work was supported in part by the Howard Hughes Medical Institute (to Y.J.) and by grant from the National Institute of Health (R35GM140892 to Y.J.). This article is subject to HHMI’s Open Access to Publications policy. HHMI lab heads have previously granted a nonexclusive CC BY 4.0 license to the public and a sublicensable license to HHMI in their research articles. Pursuant to those licenses, the author-accepted manuscript of this article can be made freely available under a CC BY 4.0 license immediately upon publication.

## Autor contributions

L.Z. and J.X. conceived the project. L.Z. prepared the samples. L.Z., Y.H., and Y.W. collected and processed the cryo-EM data and built the models. L.Z. and W.Z. designed and performed the cell-based calcium uptake assays. Y.J. supervised the research. All authors contributed to experimental design, data analysis, and manuscript preparation.

## Data and material availability

The coordinates and the Cryo-EM maps are deposited in the Protein Data Bank and Electron Microscopy Data Bank (EMDB) with accession codes: 9Z4I and EMD-73804 for rat monomeric NCLX, 9Z4J and EMD-73805 for rat trimeric NCLX. Source data are provided with this paper.

## Notes

### Competing Interest Statement

The authors have declared no competing interest.

### Summary of Updates

In this version, we have revised following parts: 1. We clarified the rationale for assigning zinc to the non-protein density at the trimerization interface in the first section of the Results. 2. We added biochemical analysis of NCLX extracted from mitochondria of NCLX-expressing cells to assess its oligomeric state in a native context. 3. We performed functional assays on mutations targeting two key structural elements of NCLX: the central calcium-binding site and the His581-mediated metal coordination site at the trimerization interface. 4. We significantly expanded the Discussion section.

