## supplementary figures for "Molecular mechanisms of mitochondrial Ca^2+^ exchanger NCLX" (from another version of this preprint: version 1, 2026-02-01)

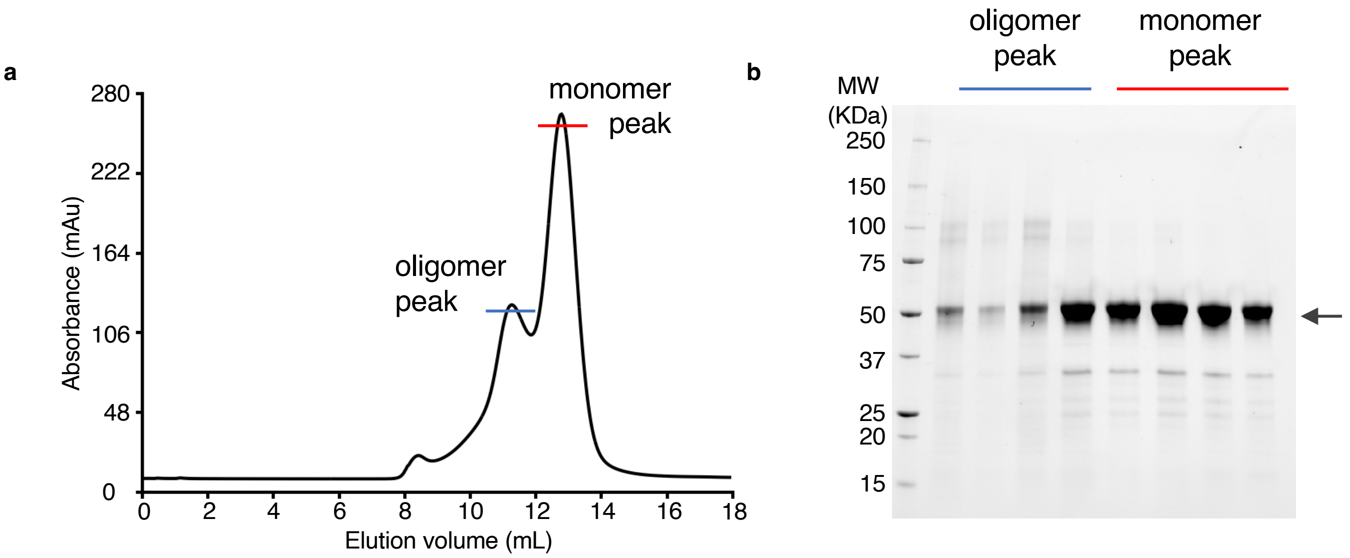


**Fig. S1: Biochemical analysis of rat NCLX. a** Size-exclusion chromatography (SEC) profile of purified NCLX showing two distinct elution peaks. **b** SDS-PAGE of fractions collected from the two SEC peaks. The arrow indicates the NCLX monomer band.

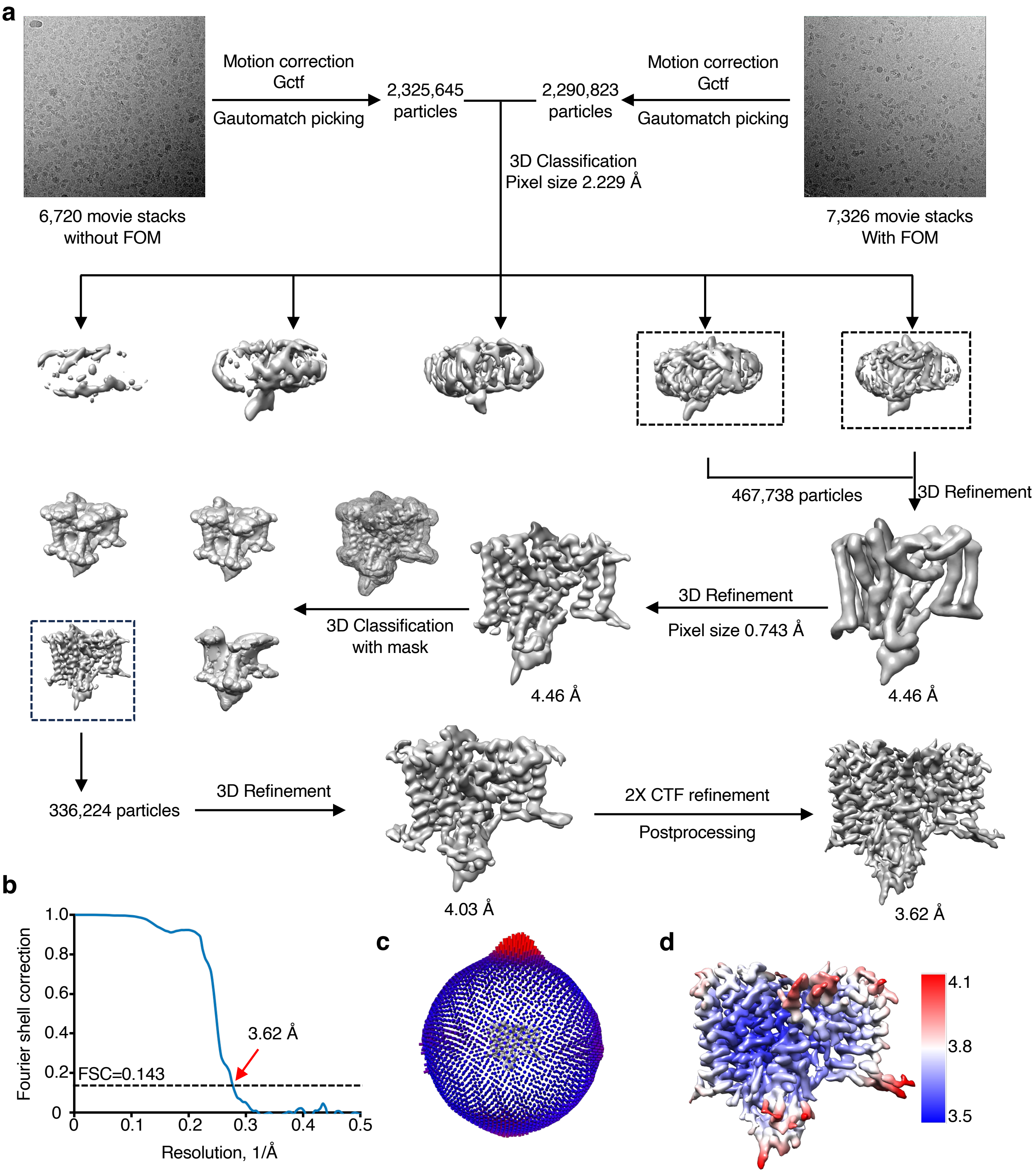


**Fig. S2: Cryo-EM data processing of monomeric NCLX**. **a** Workflow of the cryo-EM data processing. **b** Golden-standard Fourier shell correction (FSC) curve of the 3D reconstruction. **c** Angular distribution of particle projections of the 3D reconstruction. **d** Local resolution of the rat monomeric NCLX estimated using RELION.

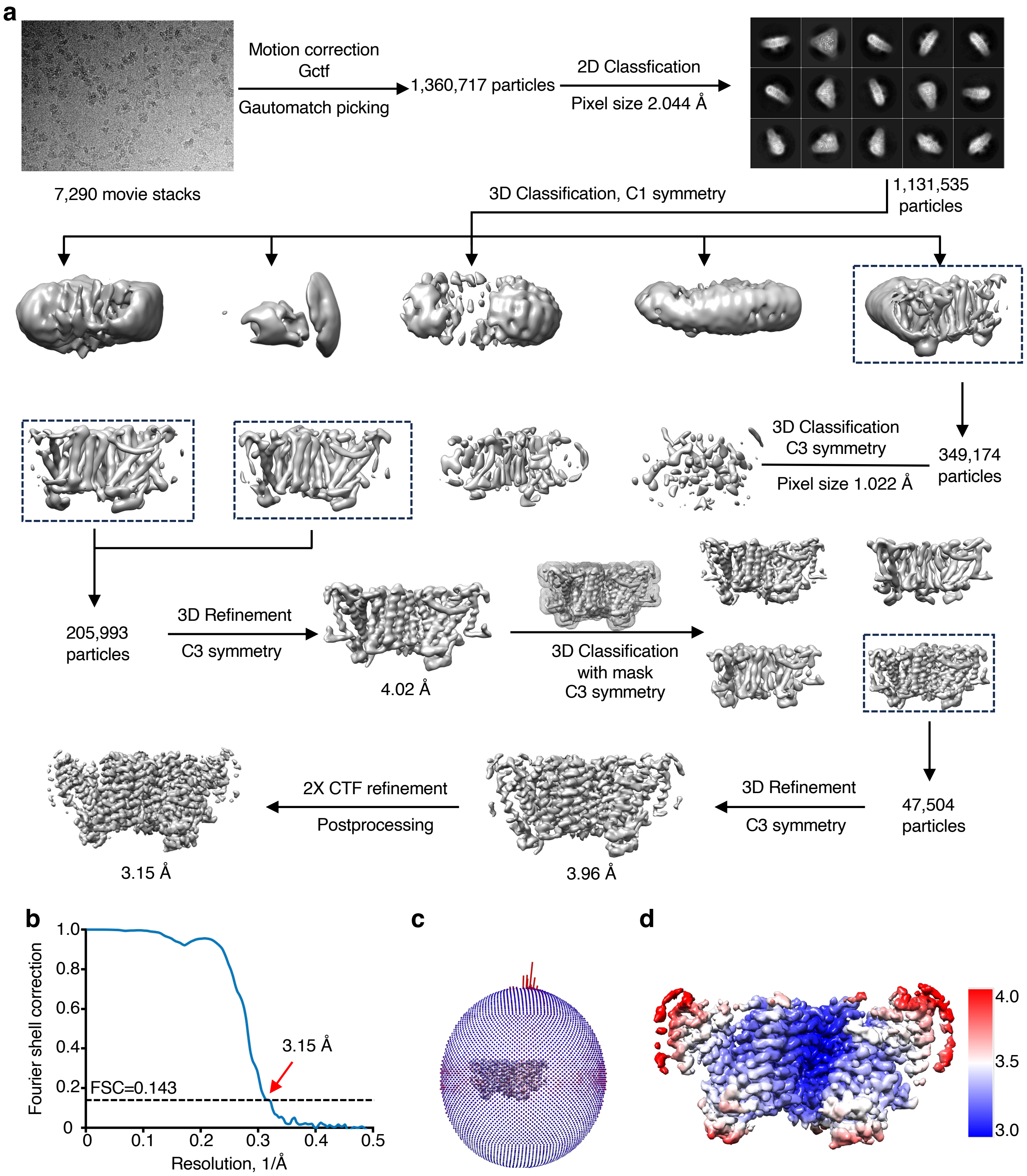


**Fig. S3: Cryo-EM data processing of trimeric NCLX**. **a** Workflow of the cryo-EM data processing. **b** Golden-standard Fourier shell correction (FSC) curve of the 3D reconstruction. **c** Angular distribution of particle projections of the 3D reconstruction. **d** Local resolution of the rat monomeric NCLX estimated using RELION.

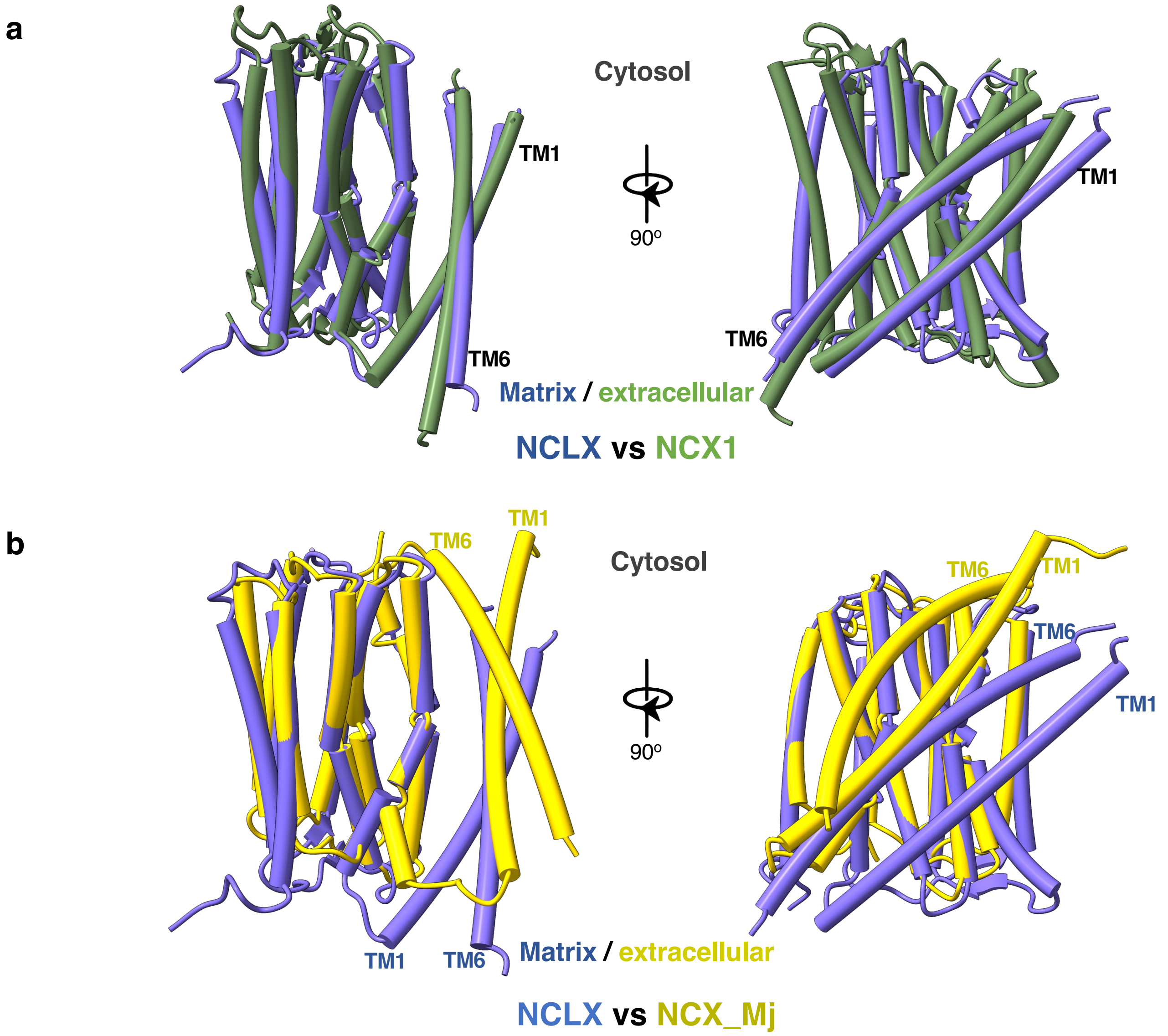


**Fig. S4: Structural comparison between NCLX and NCX. a** Comparison of the transmembrane (TM) domains between trimeric NCLX (blue) and inward-facing human NCX1 (green; PDB 8SGJ). **b** Comparison of the TM domains between trimeric NCLX (blue) and outward-facing NCX_Mj (yellow; PDB 3V5U). TMs 1 and 6 show the most pronounced structural differences between the two proteins.

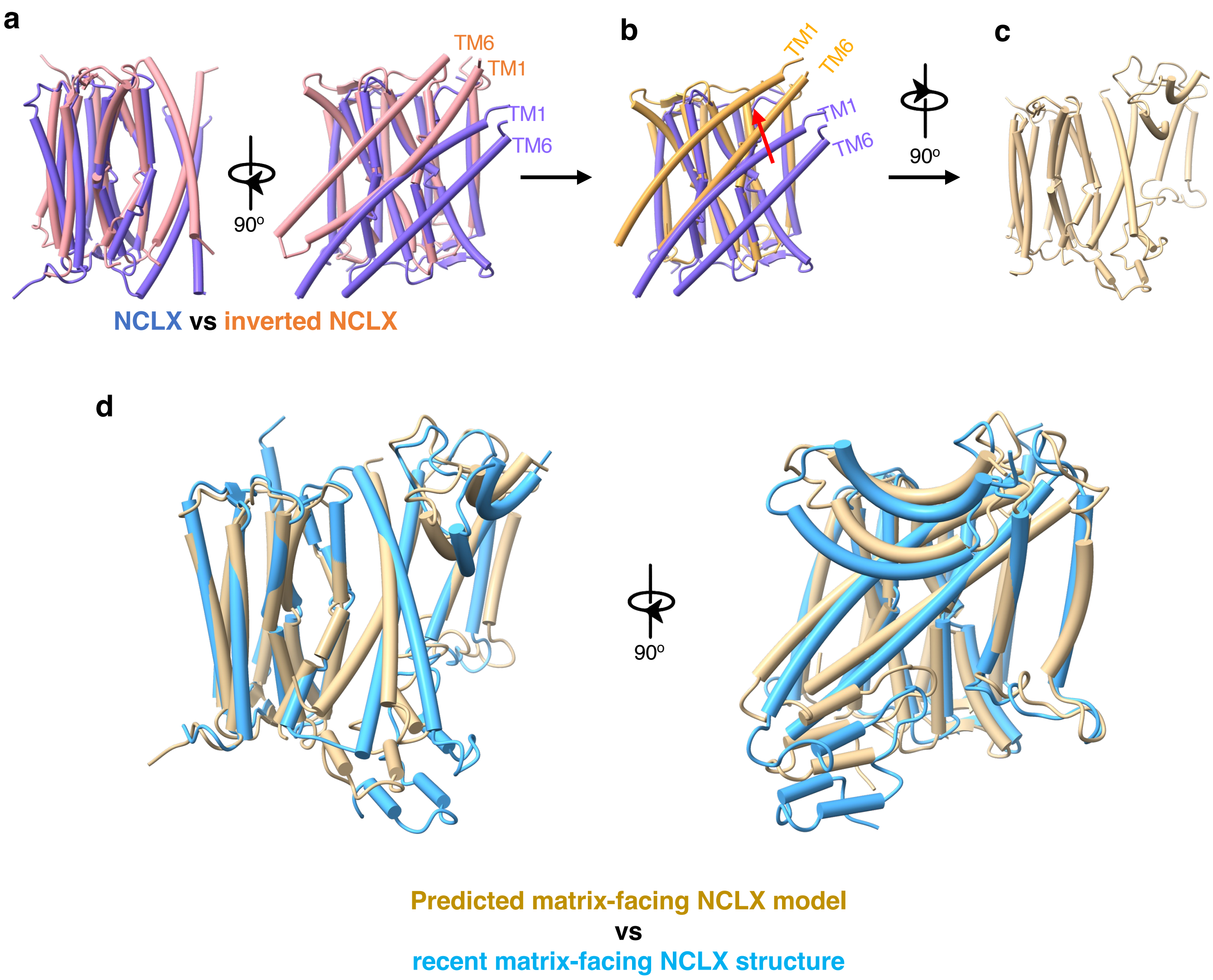


**Fig. S5: Generation of the matrix-facing NCLX model from the trimeric structure.** **a** Step 1: Superimposition of NCLX (blue) with its inverted structure (pink). **b** Step 2: TMs 1 and 6 are repositioned to the locations of inverted TMs 6 and 1, respectively, as indicated by the red arrow. **c** Step 3: The matrix-facing model is generated by moving the auxiliary components together with TMs 1 and 6 as a rigid body. **d** Comparison of the proposed matrix-facing model (light gold) with the recently reported matrix-facing NCLX structure (cyan; PDB 9PS4).

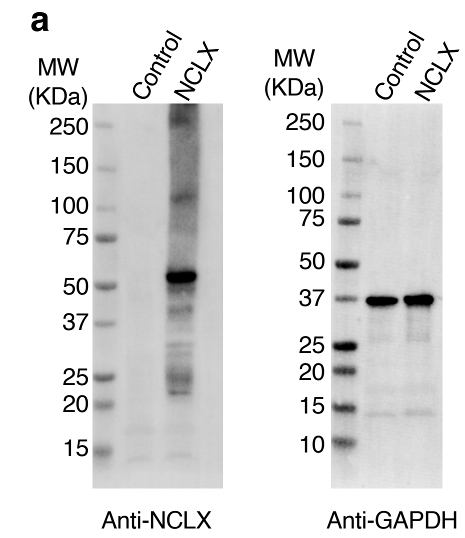


**Fig. S6. Sample preparation for $Ca^{2+}$ uptake assays.** **a** Surface expression of NCLX in HEK cells used for the $Ca^{2+}$ uptake assay. Total biotinylated surface proteins were detected using an anti-NCLX antibody. Cells transfected with empty vector served as a control. **b** Raw traces of $Ca^{2+}$ uptake in NCX1-expressing HEK cells with 150 mM extracellular $NMDG^{+}$ and varying external $[Ca^{2+}]$ (left), and corresponding traces from control HEK cells transfected with empty vector (right). Data in Fig. 5a were obtained by subtracting the control traces from the raw traces. The same procedure was applied to all $Ca^{2+}$ uptake measurements in exchanger-expressing HEK cells under various conditions shown in Fig. 5. **c** Raw sample traces of $Ca^{2+}$ uptake in NCX1-expressing HEK cells (left) and control cells (right).

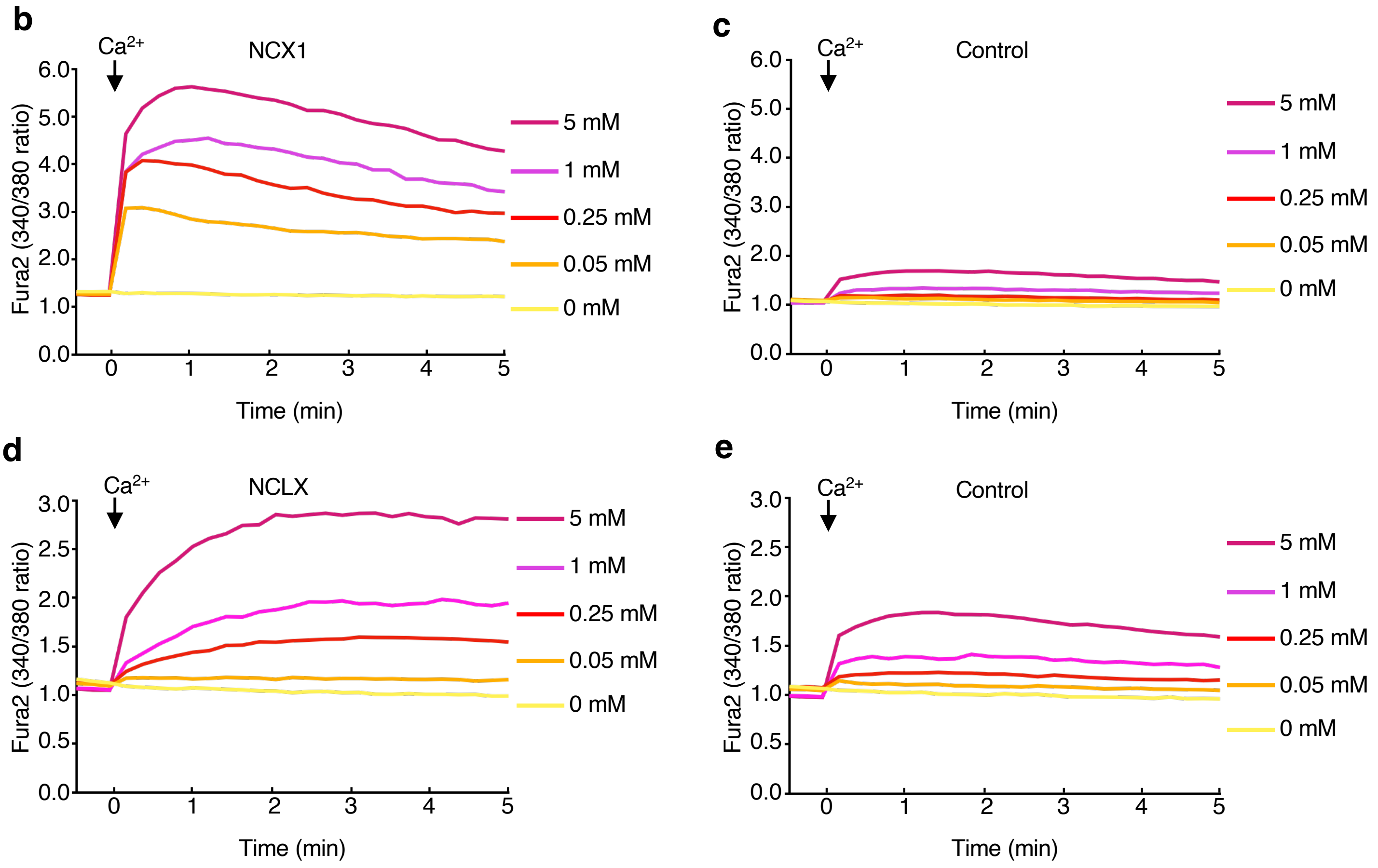
