## supplementary table for "Molecular mechanisms of mitochondrial Ca^2+^ exchanger NCLX" (from another version of this preprint: version 1, 2026-02-01)

| | NCLX_monomer (EMD-73804) (PDB 9Z4I) | NCLX_trimer (EMD-73805) (PDB 9Z4J) |
|---|---|---|
| **Data collection and processing** | | |
| Magnification | 165,000 | 130,000 |
| Voltage (kV) | 300 | 300 |
| Electron exposure ($e^-/Å^2$) | 60 | 60 |
| Defocus range (μm) | -0.8 - -2.1 | -0.8 - -2.1 |
| Pixel size (Å/pixel) | 0.743 | 0.511 |
| Symmetry imposed | C1 | C3 |
| Initial particle images (no.) | 4,616,468 | 1,360,717 |
| Final particle images (no.) | 336,224 | 47,504 |
| Map resolution (Å) | 3.62 | 3.15 |
| FSC threshold | 0.143 | 0.143 |
| **Refinement** | | |
| Initial model used (PDB code) | | |
| Model resolution (Å) | 3.65 | 3.00 |
| FSC threshold | 0.5 | 0.5 |
| Model composition | | |
| Non-hydrogen atoms | 3,748 | 11,101 |
| Protein residues | 487 | 1437 |
| Ligands | 1: Na | 3: Ca<br>1: Zn |
| B factor ($Å^2$) | | |
| Protein | 59.65 | 57.06 |
| Ligand | 30.00 | 41.28 |
| R.m.s. deviations | | |
| Bond length (Å) | 0.003 | 0.004 |
| Bond angles (°) | 0.477 | 0.585 |
| Validation | | |
| MolProbity score | 1.58 | 1.76 |
| Clashscore | 5.77 | 8.27 |
| Rotamer outliers (%) | 0.25 | 0.91 |
| Ramachandran plot | | |
| Outlier (%) | 0 | 0 |
| Allowed (%) | 3.93 | 4.44 |
| Favored (%) | 96.07 | 95.56 |
